# Trans-splicing of mRNAs links gene transcription to translational control regulated by mTOR

**DOI:** 10.1101/353979

**Authors:** Gemma Danks, Heloisa Galbiati, Martina Raasholm, Yamila N. Torres Cleuren, Eivind Valen, Pavla Navratilova, Eric M. Thompson

## Abstract

In phylogenetically diverse organisms, the 5’ ends of a subset of mRNAs are *trans*-spliced with a spliced leader (SL) RNA. The functions of SL *trans*-splicing, however, remain largely enigmatic. Here, we quantified translation genome-wide in the marine chordate, *Oikopleura dioica*, under inhibition of mTOR, a central growth regulator. Translation of *trans*-spliced TOP mRNAs was suppressed, showing that the SL sequence permits nutrient-dependent translational control of growth-related mRNAs. Under crowded, nutrient-limiting conditions, *O. dioica* continues to filter-feed, but arrests growth until favorable conditions return. Upon release from such conditions, initial recovery was independent of nutrient-responsive, *trans*-spliced genes, suggesting animal density sensing as a first trigger for resumption of development. Our results demonstrate a role for *trans*-splicing in the coordinated translational down-regulation of nutrient-responsive genes under limiting conditions.

## Introduction

*Cis*-splicing of RNA in eukaryotes removes non-coding intronic sequences from protein coding mRNAs. This is essential to their translation. In a phylogenetically disparate group of organisms (Douris, Telford, & Averof, 2010; Krchňáková, Krajčovič, & Vesteg, 2017) mRNAs also undergo trans-splicing (Hastings, 2005) where a separately transcribed RNA molecule, called a spliced leader (SL), is added to their 5’ ends. An important function of this process is to resolve polycistronic RNA transcribed from operons, allowing their translation as monocistrons. Many non-operon, monocistronic transcripts, however, are also *trans*-spliced (Blumenthal & Gleason, 2003). The function in these cases has so far remained largely enigmatic. We previously proposed the hypothesis that the SL supplies nutrient-dependent translational control motifs to *trans*-spliced mRNA (Danks et al., 2015; Danks & Thompson, 2015). TOP mRNAs, which primarily encode the protein synthesis machinery, contain a conserved 5’ Terminal OligoPyrimidine (TOP) motif that is critical (Levy, Avni, Hariharan, Perry, & Meyuhas, 1991) for translational repression during unfavourable growth conditions. This translational repression reduces the large energy expenditure associated with protein synthesis. Target of rapamycin (mTOR) (Hsieh et al., 2012; Thoreen et al., 2012), a master regulator of growth, is conserved from yeast to human and regulates the translation of mRNAs with a TOP or TOP-like motif. As part of mTORC1 (one of two complexes containing mTOR), mTOR phosphorylates and represses the translational repressor, eukaryotic translation initiation factor 4E binding protein 1 (4E-BP1). Active 4E-BP1 binds eIF4E preventing the association of eIF4E with eIF4G, which is necessary for cap-dependent translation initiation. Phosphorylated 4E-BP1 is unable to bind eIF4E and translation can proceed; mTOR thereby promotes translation and its inhibition suppresses translation.

In the urochordate, *O. dioica*, and the nematode, *C. elegans* the TOP motif is not encoded in the genome at the appropriate loci; classical TOP mRNAs in these species, therefore, lack a TOP motif. These TOP mRNAs are, however, *trans*-spliced with a pyrimidine-enriched SL sequence. The SL then forms the 5’ end where mTOR-dependent translational control is normally targeted at a TOP motif. We, therefore, concluded that these SL RNAs likely contain TOP-like mTOR-dependent translational control elements (Danks et al., 2015). This was further supported by our findings (Danks et al., 2015), and the findings of others (Zaslaver, Baugh, & Sternberg, 2011), that *trans*-spliced mRNAs are enriched for growth-related functions. In addition, we previously found that the vast majority of maternal mRNAs in the three metazoan species we examined were *trans*-spliced (Danks et al., 2015). In *O. dioica*, egg number is strongly dependent on nutrient levels (Troedsson, Bouquet, Aksnes, & Thompson, 2002) and is determined by partitioning a common cytoplasm into equally sized oocytes (Ganot, Bouquet, Kallesøe, & Thompson, 2007). This gives further indication of roles for both mTOR and *trans*-splicing in the control of maternal protein levels according to growth conditions.

Here, we tested this hypothesis by using ribosome profiling to quantify translation genome-wide in female *O. dioica* treated with the mTOR inhibitor Torin 1 (Thoreen et al., 2012, 2009). The mTOR-regulated translatome was conserved between *O. dioica* and other species. Moreover, classical TOP mRNAs that possess a 5’ *trans*-spliced SL sequence rather than an encoded TOP motif were nevertheless subject to translational control via the mTOR pathway in *O. dioica*. These results suggest that *trans*-splicing may play a key role in the coordinated nutrient-dependent translational regulation of growth-related genes

Under conditions of nutrient depletion and high animal density, *O. dioica* enters a developmental growth-arrested state (stasis), prior to the onset of meiosis, and mTOR activity is down-regulated (Subramaniam, Campsteijn, & Thompson, 2014) indicating that the translation of TOP mRNAs is suppressed. Once conditions become more favourable the animals recover and resume normal development Danks et al. (2015); Subramaniam et al. (2014). In the absence of food, the nematode *C. elegans* also enters a state of developmental growth arrest (L1 diapause). When food becomes available, animals resume development. In *C. elegans*, transcription of operons is preferentially up-regulated during recovery from growth arrest (Zaslaver et al., 2011). In *O. dioica*, however, it is instead non-*trans*-spliced monocistrons that are transcriptionally up-regulated during recovery from growth arrest (Danks et al., 2015). We previously hypothesized, therefore, that during recovery from growth arrest *O. dioica* up-regulates TOP mRNAs and other *trans*-spliced transcripts via translational control, rather than transcriptional control, as a faster initial response (Danks et al., 2015). Here, we tested this hypothesis using ribosome profiling on *O. dioica* during growth arrest and recovery and found that, as with transcription, the translation of *trans*-spliced genes, including mTOR-targeted *trans*-spliced TOP mRNAs, were not preferentially up-regulated during recovery. This suggests that the primary, first response during recovery is not mediated by coordinated, enhanced translation of TOP mRNAs in *O. dioica.*

Our data support the hypothesis that *trans*-splicing in metazoans plays a key role in nutrient-dependent translational control and evolved as a result of the advantage it provides in the allocation of maternal resources, during vitellogenesis, according to nutrient levels, rather than in growth arrest recovery.

## Results

We first profiled translation genome-wide in day 6 female *O. dioica.* At this stage the bulk of the animal’s mass is from a single-celled coenocyst (Ganot, Bouquet, et al., 2007; Ganot, Kallesøe, & Thompson, 2007; Ganot, Moosmann-Schulmeister, & Thompson, 2008) within the ovary, the transcriptional output of which is enriched for *trans*-spliced transcripts (Danks et al., 2015). Second, we confirmed that exposing female *O. dioica* to the mTOR inhibitor Torin 1, in seawater, resulted in the expected absence of phosphorylated 4E-BP1 (Figure 1A and Figure S1). Phosphorylated 4E-BP1 was absent after 1.5 h of treatment, similar to what was observed in mouse embryonic fibroblast (MEF) cells (Thoreen et al., 2012). Polysome profiles confirmed the down-regulation of translation as indicated by reduced polysome peaks (Figure S2). We then measured the effect of mTOR inhibition on the translation of individual mRNAs by quantifying and precisely mapping ribosome protected RNA fragments (RPFs) using ribosome profiling with deep sequencing (Ingolia, Brar, Rouskin, McGeachy, & Weissman, 2012). In parallel, we sequenced total RNA in order to normalise RPFs to the abundance of mRNA transcripts (RNA). Sequencing generated 57.9M (vehicle control: DMSO) and 42.8M (Torin 1) total RNA exon-mapped reads and 24.1M (DMSO) and 1.4M (Torin 1) RPF exon-mapped reads, across three biological replicates (Table S1). By excluding genes with low read counts we were able to confidently assess the translational efficiency of 14,574 expressed genes out of 17,212 in the *O.dioica* reference genome.

### The mTOR-regulated translatome is conserved between *O. dioica* and vertebrates and enriched for *trans*-spliced TOP mRNAs

We normalised RPF read counts to RNA read counts to quantify translational efficiency (TE). This allows the detection of mRNAs with unusually high or low ribosome density given their transcript abundance. We detected 762 genes with mRNAs that had significantly reduced translational efficiencies when mTOR was inhibited (Figure 1B and Table S2). These represent the main targets of mTOR-mediated translational control in female *O. dioica.* Gene ontology (GO) analysis revealed that these were enriched for functions known to be regulated by TOR signalling, including translation and translation elongation, mitotic spindle elongation, fatty acid metabolism and TOR signalling (Figure S3). Importantly, these include known TOP mRNAs: 60/127 expressed *O. dioica* ribosomal protein mRNAs and 59/90 mRNAs with orthologs to known human TOP mRNAs were significantly down-regulated (Figure 1C,E and Table S3). As found in mammalian cells, histone mRNAs were amongst those resistant to Torin 1 (Figure 1C and Table S2). Together, our data show that the targets of mTOR regulation are conserved between *O. dioica* and vertebrates.

### Translation of *trans*-spliced TOP mRNAs is regulated by mTOR

The TOP motif in vertebrate canonical TOP mRNAs (ribosomal proteins and other members of the translational apparatus) is highly conserved and required for growth-dependent translational control via mTOR signalling (Levy et al., 1991). A 5’ TOP motif is also enriched in ribosomal protein mRNAs in the ascidian *Ciona intestinalis* (*Danks et al., 2015*; *Yokomori et al., 2016*). The canonical TOP motif begins with a cytosine and is followed by a stretch of 4-14 pyrimidines (Meyuhas, 2001). It was recently shown in MEFs that mTOR regulates a broader spectrum of mRNAs (Thoreen et al., 2012). These are enriched for the presence of a TOP-like pyrimidine-enriched motif (a stretch of at least 5 pyrimidines within 4 nucleotides of the transcription start site (TSS)) (Thoreen et al., 2012). The majority of established TOP mRNAs, including those discovered recently, are *trans*-spliced in *O. dioica* (*Danks et al., 2015*). These include 103 out of 129 (80%) ribosomal proteins, 33 out of 40 eukaryotic translation initiation factors (including 4 out of 5 that are known TOP mRNAs), eukaryotic elongation factor 1A, eukaryotic elongation factor 2, translationally controlled tumour protein (TCTP), vimentin and rack1 (Table S3). All these TOP mRNAs receive the 40 nt spliced leader (SL) RNA sequence at their 5’ ends. The 5’ end of this SL sequence (Ganot, Kallesøe, Reinhardt, Chourrout, & Thompson, 2004) (ACTCATCCCATTTTTGAGTCCGATTTCGATTGTCTAACAG) is pyrimidine-enriched (12 out of the first 15 nucleotides are pyrimidines), although it starts with an adenine and is interrupted by several purines. This suggests that the 5’ end of the spliced-leader can function as a TOP motif in the mTOR-mediated regulation of translation. Our data confirmed that the translation of *trans*-spliced TOP mRNAs was suppressed upon the inhibition of mTOR: 51 out of the 60 *O. dioica* ribosomal protein mRNAs with translation significantly repressed by mTOR inhibition are *trans*-spliced (Table S3).

**Figure 1.**
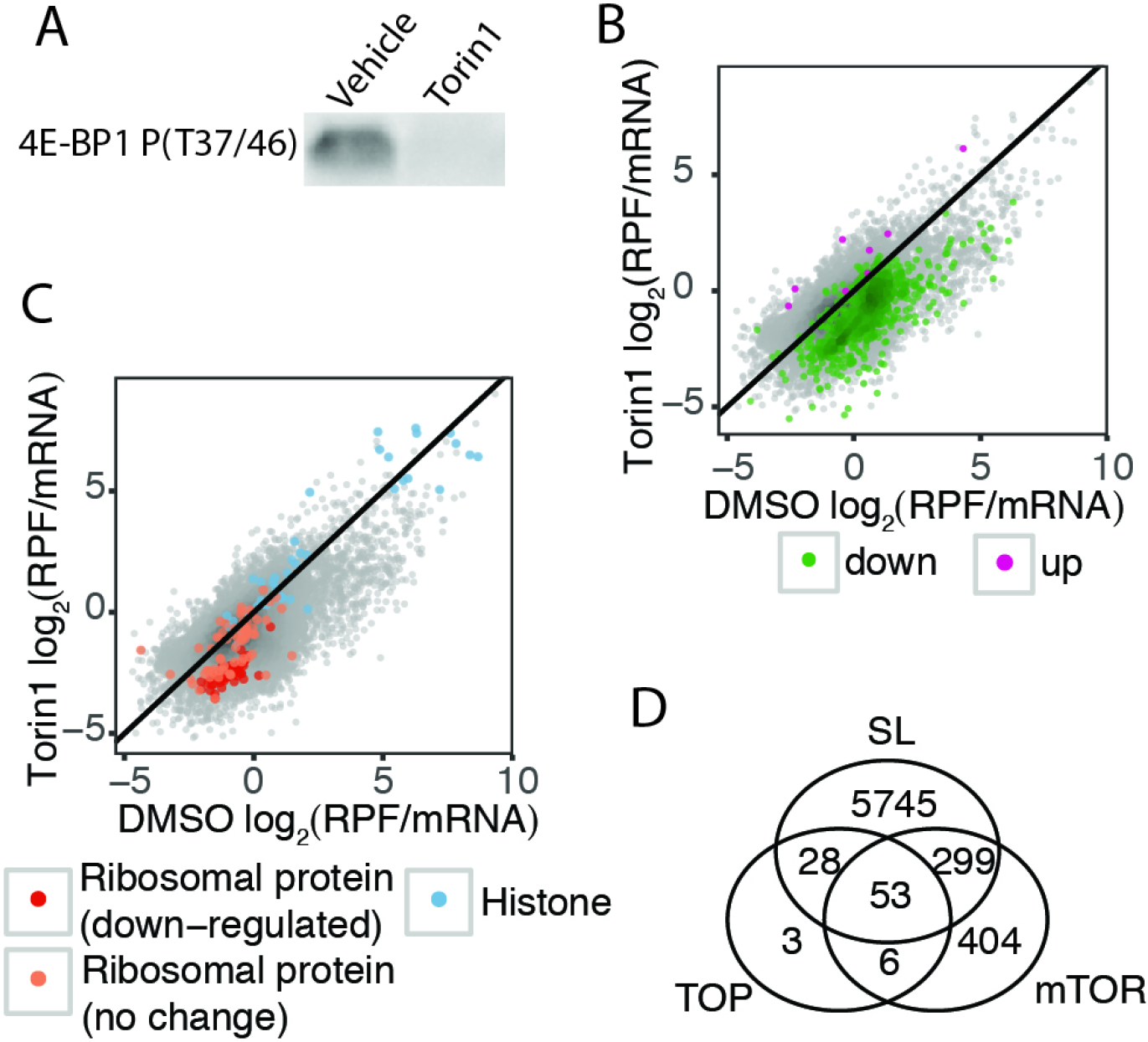
The mTOR-regulated translatome of O. *dioica*. (**A**) Adult animals were exposed to the mTOR inhibitor, Torin 1 (1 μM), or DMSO (vehicle control) in seawater for 1.5 h and female animals were collected. Phosphorylated 4E-BP1 was detected in DMSO but not in Torin 1 treated animals, confirming that mTOR was inhibited in Torin 1-treated animals (one out of three replicates shown; total protein used as loading control and normalisation of band intensity; see Figure S1D for full blot). (**B**) Median translational efficiency (RPF/mRNA= ribosome protected fragment density/mRNA density) of mRNAs from 3 replicates for Torin 1- and DMSO-treated animals with transcripts identified as having significantly up-or down-regulated translation highlighted. (**C**) Translational efficiencies shown as in (**B**) with known Torin 1-resistant (histone mRNAs) and Torin 1-sensitive (ribosomal protein mRNAs) gene categories highlighted to show that targets of mTOR-mediated translational control are conserved in *O. dioica*. (**D**) Intersections of orthologs of known TOP mRNAs (TOP), mTOR-regulated mRNAs (mTOR) and mRNAs that are *trans*-spliced (SL).

### *Trans*-spliced transcripts dominate the primary translational response to mTOR inhibition

*Trans*-splicing of mRNAs is not limited to TOP mRNAs but is associated with 39% of all *O. dioica* genes, a subset that is enriched for a wider array of functions related to growth (Danks et al., 2015). Of the female-expressed genes that we analysed, 43% are *trans*-spliced. Since the translation of *trans*-spliced TOP mRNAs is mediated via mTOR in *O. dioica*, it follows that all *trans*-spliced mRNAs are potential targets for growth-dependent translational control, although this is not significantly more than expected given the frequency of *trans*-splicing. Interestingly, however, we found that mRNAs that were suppressed only at the translation level, and not at the transcription level, were enriched for *trans*-spliced transcripts (56% of mRNAs with translation-only suppression are *trans*-spliced compared to 34% of those with both translational and transcriptional responses to mTOR inhibition; Fisher’s exact test P-value = 2.97 × 10^−9^) (Figure 2B). This indicates that *trans*-spliced transcripts dominate the primary translational response to mTOR inhibition and *non-trans*-spliced transcripts constitute a longer-term, secondary response involving additional, slower transcriptional adjustment of gene expression. Indeed, GO analysis of these subsets revealed that genes with a transcriptional response to mTOR inhibition were enriched for functions related to proteolysis and muscle contraction (Figure 2A), the latter being characteristic of genes with transcription down-regulated during growth arrest in *O. dioica* (Danks et al., 2015).

**Figure 2:**
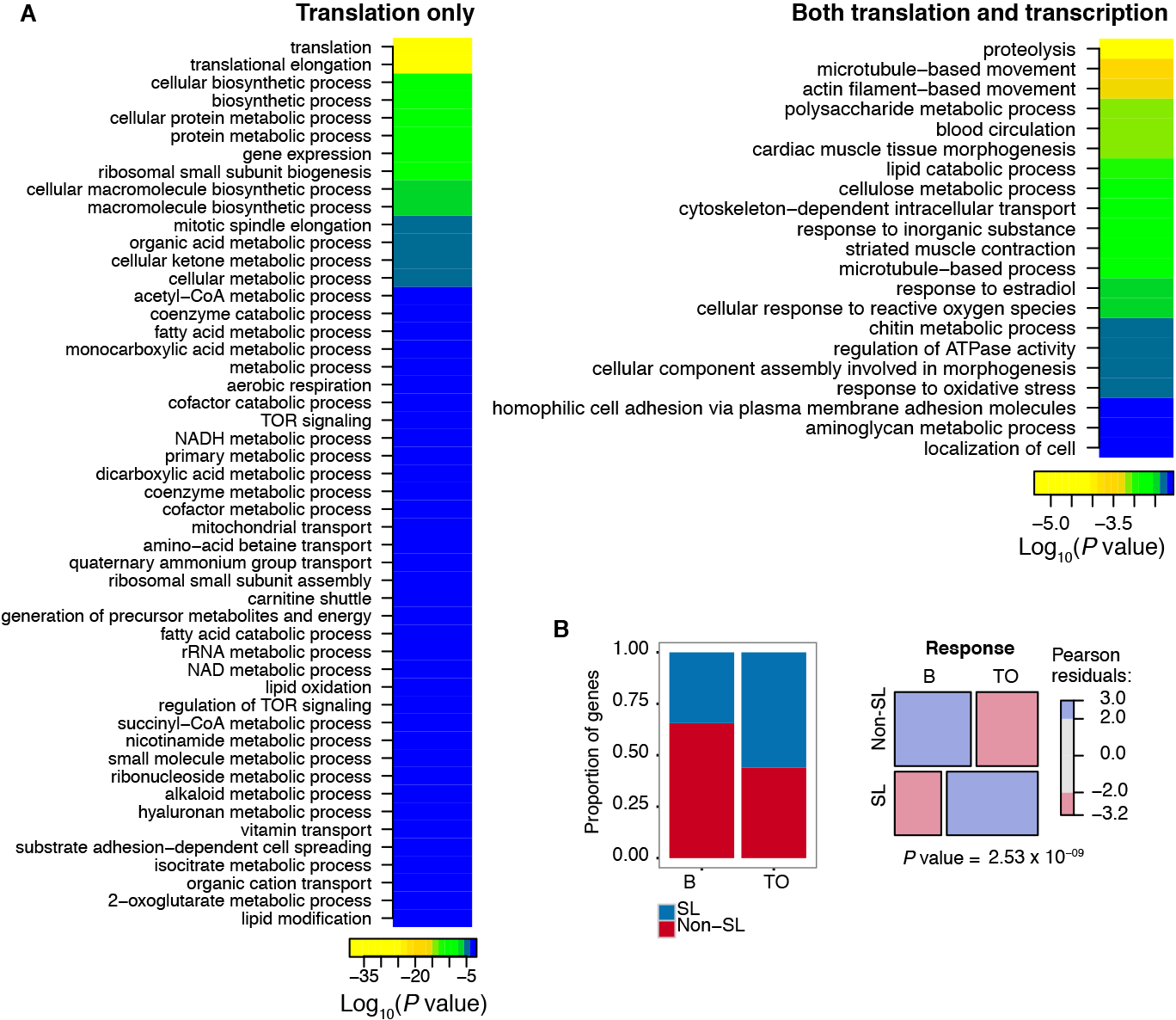
Translational and transcriptional responses to mTOR inhibition. (**A**) GO analysis of genes with significantly down-regulated translation upon inhibition of mTOR where the response is translational only (left), indicative of a primary response, or both translational and transcriptional (right), indicative of a secondary longer-term response (transcriptional-only comprised only 7 genes and were not analysed further) (**B**) Bar chart (left) shows the proportion of genes with significant down-regulation of translation that are *trans*-spliced (SL) for each response category in (A): TO = translation only; B = both translational and transcriptional response. Mosaic plot (right) visualises the Pearson residuals from a corresponding χ^2^ test.

### Oocyte-stocked mRNAs are *trans*-spliced and translationally dormant

Surprisingly, given the clear regulation of translation of *trans*-spliced TOP mRNAs, *trans*-spliced genes in female *O. dioica* were, on average, more resistant to mTOR inhibition (mean log_2_ (ΔTE) = −0.28) than genes that were not trans-spliced (mean log_2_ (ΔTE) = −0.74) (Welch two sample t-test: t = 24.484, df = 14384, P-value < 2.2 × 10^−16^) (Figure 3A). *Trans*-spliced transcripts that were not suppressed had significantly lower mean translational efficiencies (mean TE = 1.12), under control conditions, than transcripts that were not *trans*-spliced (mean TE = 2.76) (Welch two sample t-test: t = -9.6612, df = 11967, P-value < 2.2 × 10^−16^) (Figure 3B) and a significantly higher mRNA abundance (Welch two sample t-test: t = 74.861, df = 12970, P-value < 2.2 × 10^−16^) (Figure 3C). This low level of translation under normal conditions may explain why these transcripts are insensitive to translational suppression via mTOR inhibition. The high abundance but low translation of these mRNAs suggests that they are sequestered: most likely in arrested oocytes, which contain a large fraction of the total RNA pool in this stage of the female *O. dioica* lifecycle and where the majority of transcripts are *trans*-spliced (Danks et al., 2015). We performed a fluorescent detection of nascent protein synthesis as well as polysome profiling to confirm that mRNAs in oocytes are dormant (Figure 3F,G). The majority of our weakly-translated, Torin 1-resistant, *trans*-spliced transcripts therefore likely represent dormant maternal mRNAs stocked in oocytes. We used both tiling microarray (Danks et al., 2013) and cap analysis of gene expression (CAGE) (Danks, Navratilova, Lenhard, & Thompson, 2018) data from *O. dioica* oocytes to determine the set of oocyte-stocked mRNAs. As expected, we found that the translational efficiency of oocyte transcripts in control animals was significantly lower than that of non-oocyte transcripts (mean oocyte log_2_ (TE) = −0.80; mean non-oocyte log_2_ (TE) = 0.45; Welch two-sample t-test: t = −57.494, df = 14002, P-value < 2.2 × 10^−16^) (Figure 3D, Figure S4), and the effect of Torin 1 was significantly reduced (mean oocyte log_2_ (Δ) = -0.17; mean non-oocyte log_2_ (Δ) = -0.97; Welch two-sample t-test: t = 43.54, df = 12832, P-value < 2.2 × 10^−16^) (Figure 3E). Importantly, we found that 80% (4639/5772) of Torin 1-resistant *trans*-spliced transcripts were present in the oocyte. When we removed oocyte transcripts from our analysis we found that transcripts with suppressed translation upon mTOR inhibition were enriched for those *trans*-spliced with the SL (28.6% of down-regulated genes are *trans*-spliced compared to 17.5% of unaffected genes; Fisher’s exact test P-value = 1.26 × 10^−7^). This is despite excluding most TOP mRNAs, which have transcripts present in the oocyte. We obtained similar results when excluding all transcripts with low levels of translation (DMSO log_2_ (TE) < 1) under control conditions (36.6% of down-regulated genes were *trans*-spliced compared to 22.9% of unaffected genes; Fisher’s exact test P-value = 8.4 × 10^−11^).

### A TOP motif is not encoded in the genes of mTOR-regulated transcripts

We wanted to determine whether or not a TOP-like motif is present at the 5’ ends of translation-suppressed transcripts that were not *trans*-spliced. We obtained transcription start sites (TSSs) at bp-resolution in female animals from CAGE data (Danks et al., 2018) and examined the 5’ sequences of all expressed transcripts. Out of 2,772 robustly expressed, *non-trans*-spliced transcripts, only 4 had a canonical TOP motif and only 66 had 5’ pyridine-enrichment comparable to the SL sequence. A more relaxed definition of a TOP-like motif (at least 5 pyrimidines within 4 nucleotides of a TSS) (Thoreen et al., 2012) was also only present at a low frequency (0.076); lower than in mammalian cells (0.16) (Thoreen et al., 2012). We found no significant enrichment of this motif at the 5’ ends of transcripts that had suppressed translation upon mTOR inhibition in *O. dioica*. Since the SL has a stretch of 5 pyrimidines further downstream we also relaxed the definition of this motif further but still found no significant enrichment in suppressed transcripts. This indicates that these transcripts are indirect targets of translational suppression resulting from a global down-regulation of translation. Supporting this hypothesis, a GO term analysis of mTOR-regulated transcripts lacking a spliced leader revealed an enrichment of functions related to autophagy (proteolysis) and lipid catabolism whereas those that are *trans*-spliced are enriched for known TOP mRNA functions related to protein synthesis.

### *Trans*-spliced transcripts in *C. elegans* are under growth-dependent translational control

We next sought to identify *trans*-spliced TOP mRNAs that are under mTOR regulation in another metazoan species. *C. elegans trans*-splices 70% of its mRNAs to one of two pyrimidine-enriched spliced leaders (Allen,Hillier, Waterston, & Blumenthal, 2011; Blumenthal & Gleason, 2003). SL1 is associated with monocistrons and the first gene in an operon and SL2 is associated with downstream operon genes. Included amongst these are all but one ribosomal protein genes (TOP mRNAs), which are mostly *trans*-spliced with SL1 (Danks et al., 2015). A genome-wide study of translation during L1 diapause exit identified ribosomal protein mRNAs as transcripts with the highest translational up-regulation (Stadler & Fire, 2013). While no mention of the association of these transcripts with *trans*-splicing was made in this study, the data nevertheless clearly show that *trans*-spliced ribosomal protein (TOP) mRNAs are targets of nutrient-dependent translational-control. Furthermore, a recent study showed that *trans*-splicing in *C.elegans* enhances translational efficiency (Yang et al., 2017). In order to establish whether or not a relationship exists between the presence of SL1 and/or SL2 at the 5’ end of an mRNA and its translational control during recovery from growth arrest we re-analysed existing ribosome profiling and mRNA-seq data from L1 diapause exit (Stadler & Fire, 2013) together with existing data mapping *trans*-splice sites genome-wide in *C. elegans* (Allen et al., 2011). We used a total of 10,362 genes that could be tested for differential translational regulation and assigned a *trans*-splicing category with high confidence. Amongst these, we found a strong relationship between the presence of a 5’ spliced leader and translational control during L1 diapause exit in response to food availability (χ^2^ =711.45, df = 4, *P* value < 2.2 × 10-16) (Figure S5). Amongst transcripts with up-regulated translation, 54% (786/1460) are *trans*-spliced to SL1 and 18% (260/1460) are trans-spliced to SL2, while 414 (28%) lack a 5’ spliced leader (Figure S5). This constitutes an enrichment of *trans*-spliced transcripts compared to unaffected transcripts (56% of which lack a 5’ spliced leader). These results show that *trans*-spliced TOP mRNAs in *C. elegans* are also under nutrient-dependent translational control, indicating that the spliced leaders in *C. elegans* are targets of mTOR.

**Figure 3.**
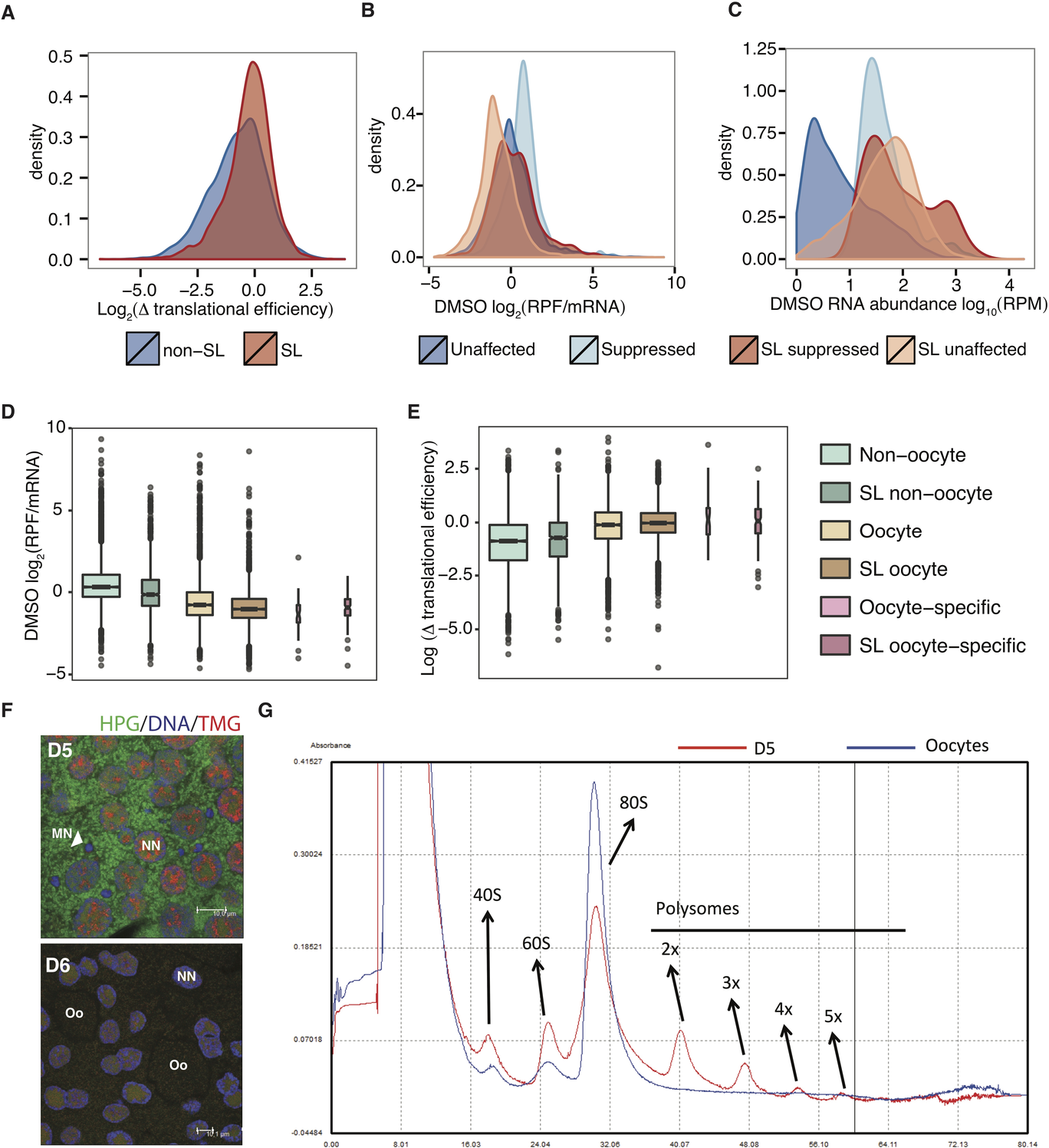
Abundant maternal mRNAs stocked in the oocyte are *trans*-spliced, translationally dormant and resistant to mTOR-inhibition. (**A**) Distribution of changes in translational efficiency in response to Torin 1 for all transcripts with and without a 5’ spliced leader (SL) sequence. (**B**) Distribution of translational efficiencies (ribosome density normalised to mRNA abundance; RPF = ribosome protected fragments) in control animals with transcripts categorised according to the presence of the 5’ spliced leader and their response to mTOR inhibition (suppressed or unaffected). (**C**) Distribution of mRNA abundances (RPM = reads per million) in control animals with transcripts categorised as in the colour legends under **B** and **C**. (**D**) Distribution of translational efficiencies in control animals with transcripts categorised by the presence of the 5’ spliced leader and whether or not they are present in oocytes (detected by cap analysis of gene expression (CAGE) or tiling microarray from oocyte samples). (**E**) Distribution of Torin 1-induced changes in translational efficiency for transcripts categorised as in (**D**). (**F**) The cytoplasm of the coenocyst in day 5 (D5) female gonads, pre-oocyte formation, has a high level of translational activity (as indicated by the intensity of HPG). Oocytes (Oo) that have formed by late day 6 (D6), however, are translationally dormant. (**G**) RNA from D5 animals have higher levels of polysome occupancy compared to RNA from oocytes, where there are no visible polysome peaks.

### Exit from growth arrest in *O. dioica* is not dependent on mTOR

Having established that *trans*-spliced transcripts in female *O. dioica* are targets of mTOR-regulated translational control and that translation of *trans*-spliced TOP mRNAs are up-regulated during recovery from growth arrest in *C. elegans*, we next wanted to assess the translational regulation of *trans*-spliced transcripts during recovery from growth arrest in *O. dioica.* We previously proposed that translational control, rather than transriptional control, may up-regulate *trans*-spliced growth-related genes during recovery (Danks et al., 2015). We performed ribosome profiling on *O. dioica* during growth arrest (stasis, high animal density) and recovery from growth arrest (release into normal animal density). Sequencing generated 27.0M (stasis) and 38.6M (release) total RNA exon-mapped reads and 1.8M (stasis) and 1.5M (release) RPF exon-mapped reads, across two biological replicates. We detected 1601 genes with significantly up-regulated transcription and 638 with significantly down-regulated transcription during release from stasis. We confirmed our previous observations (Danks et al., 2015) that genes that are transcriptionally up-regulated are enriched for muscle-related GO terms and that *trans*-splicing is under-represented (Figure S6). We then analysed differential translational efficiency and found 1382 genes with significantly up-regulated translational efficiency upon release from stasis and only 28 significantly down-regulated (Figure 4). Surprisingly, we found that only 8/129 ribosomal protein mRNAs were up-regulated (Figure 4b). *Trans*-spliced transcripts were not over-represented in the set of up-regulated genes and the mean change in translational efficiency was not significantly different between trans-spliced and non-trans-spliced transcripts (t-test: t = −0.32652, df = 13368, *P* value = 0.744). GO terms that were over-represented in the set of genes with up-regulated translational efficiencies included terms related to muscle contraction, hormone regulation and the cell cycle (Figure S7), rather than terms typical of the mTOR-dependent translatome identified above. These results show that up-regulation of nutrient-dependent growth-related genes is not the initial response to release from growth arrest in *O. dioica.* Supporting this, replication tracing by EdU incorporation showed that endocycling, which is suppressed during growth arrest, resumed in released animals regardless of whether or not food was available (Figure 4C-G).

## Discussion

We have shown that *trans*-splicing of a spliced leader sequence to the 5’ ends of mRNAs is associated with growth-dependent translational control in two different metazoans. Inhibiting mTOR in *O. dioica* revealed a typical mTOR-dependent translatome with classical TOP mRNAs as primary targets. These TOP mRNAs do not contain the highly conserved TOP motif, as in other species, but are instead *trans*-spliced with a pyrimidine-enriched TOP-like spliced leader sequence that can perform the same role as a TOP motif in permitting nutrient-dependent translational control of mRNAs via mTOR (Figure 5). Nutrient-induced recovery from L1 arrest in *C. elegans* involves the translational up-regulation of mTOR-regulated transcripts that are *trans*-spliced. The potential survival advantage of an association between mTOR and the spliced leader may be a key driving force in the evolution (or retention) of *trans*-splicing. The *trans*-splicing of translational control motifs also permits the rapid evolution of new targets for mTOR-regulation, given that all that is required is an unpaired acceptor site for *trans*-splicing of the spliced leader, rather than the evolution of the complete TOP motif. In addition, the regulation of *trans*-splicing itself allows for the switching on or off of mTOR targets at different stages of the life cycle. This may be achieved through the use of alternative start sites that either include or exclude the *trans*-splice site. Supporting this, a male-specific promoter motif recently identified in *O. dioica (Danks et al., 2018)* causes the exclusion of *trans*-splice sites, and consequently the lack of a spliced leader on resulting transcripts, in many cases. Given the strong association of *trans*-splicing with maternal mRNA, mTOR regulation via the spliced leader is likely an underlying mechanism for the adjustment of egg numbers according to nutrient levels in *O. dioica (Troedsson et al., 2002).* This implicates *trans*-splicing as a master, molecular-level mediator of population levels in this abundant zooplankton.

**Figure 4.**
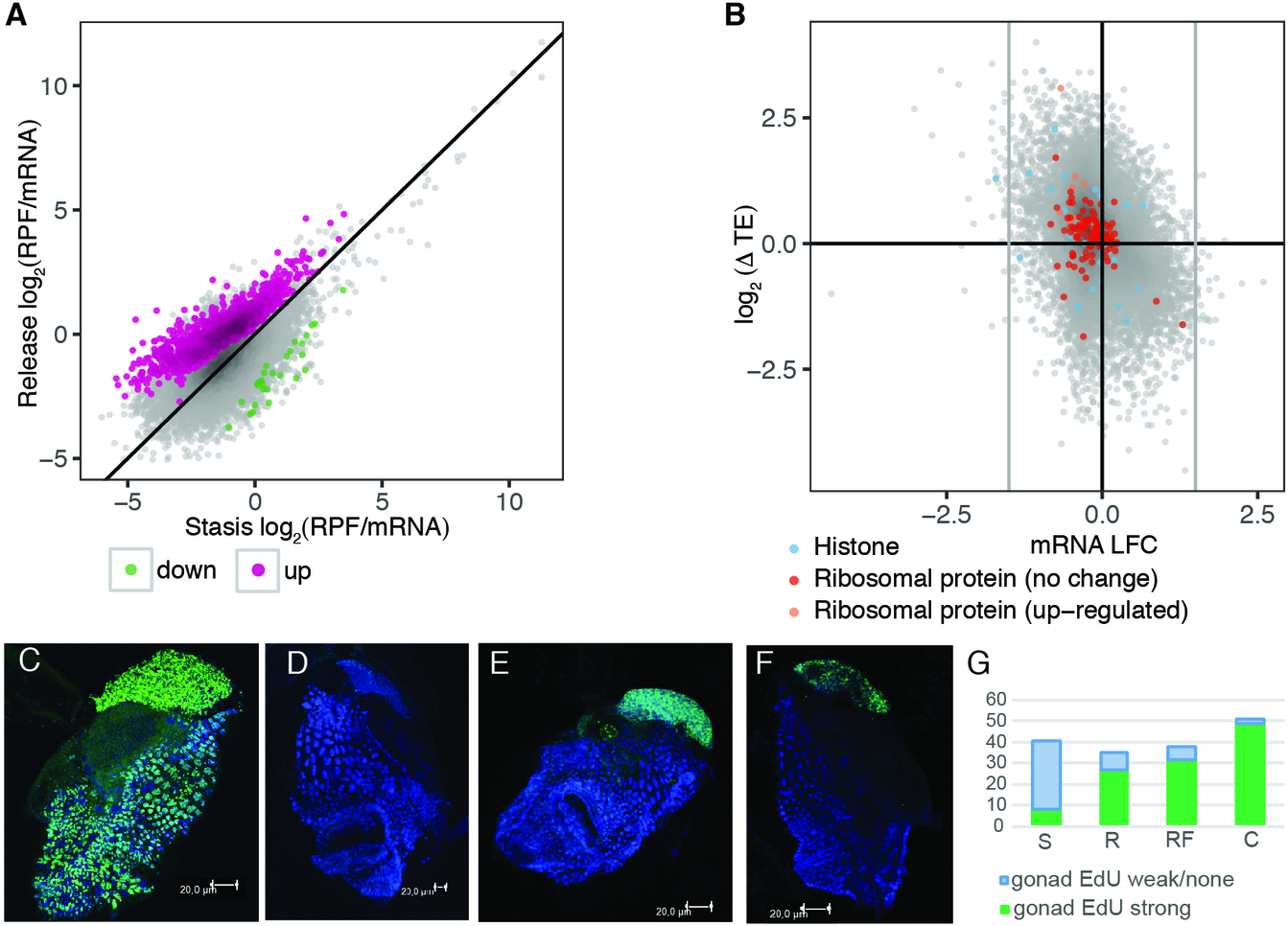
The growth arrest recovery translatome of *O. dioica.* Animals were cultured under dense conditions before being released by dilution in the presence of food. (**A**) Median translational efficiency (RPF/mRNA= ribosome protected fragment density/mRNA density) of mRNAs from 2 replicates for growth arrested and released animals with transcripts identified as having significantly up-or down-regulated translation highlighted. (**B**) Degree of change in translational efficiency upon release from stasis (y-axis) against log fold change in mRNA abundance (x-axis) with known mTOR-independent (histone mRNAs) and mTOR target (ribosomal protein mRNAs) gene categories highlighted. The majority of mTOR targets are not up-regulated upon release from stasis. (**C-G**) EdU incorporation (DNA replication) was restored in the germline 12 h after animals were released from the crowded conditions of growth arrest independent of food supply confirming that a change in density rather than increased food availability is the primary trigger for exiting a growth-arrested state in *O. dioica:* (**C**) growth arrest; (**D**) normal day 3; (**E**) release without food; (**F**) release with food; (**G**) proportion of animals showing extensive DNA synthesis under all four conditions (S = stasis; R = release without food; RF = release with food; C = normal day 3 control).

**Figure 5.**
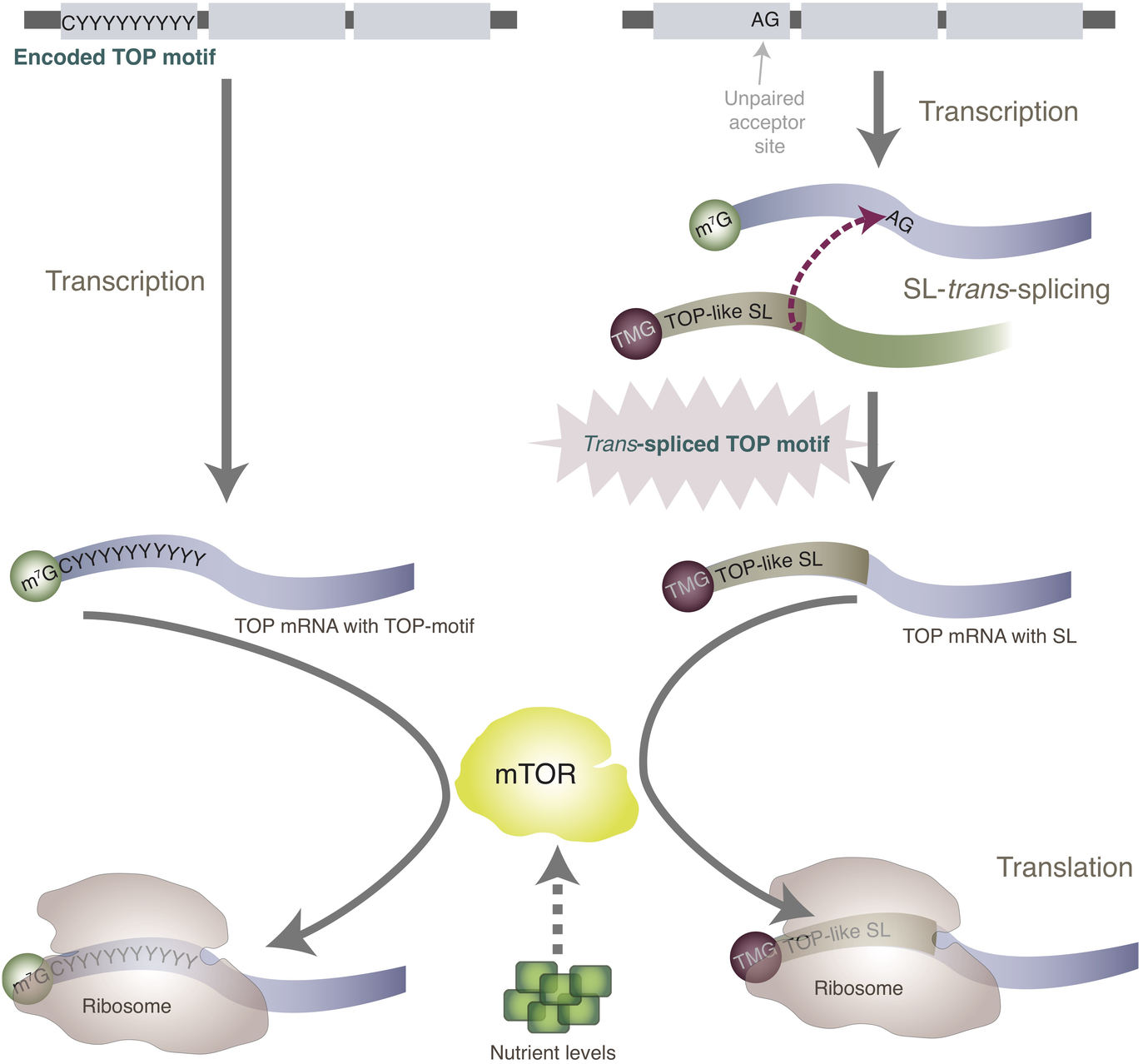
Translational control of TOP mRNAs. Schematic compares the translational control of TOP mRNAs in animal genomes where a TOP motif is encoded at gene loci (left) and in animals where a TOP-like sequence is added post-transcriptionally via *trans*-splicing (right). The nutrient-dependent translation of classical TOP mRNAs (predominantly ribosomal proteins) in mammalian cells is controlled via mTOR signalling and requires a 5’ Terminal Oligopyrimidine Tract (TOP motif). In *O. dioica* and *C. elegans*, the TOP motif is not encoded in the 5’ UTRs of many of such genes, but instead, these mRNAs are *trans*-spliced with a spliced leader (SL) that has a pyrimidine-enriched 5’ end. We confirmed that the translation of these *trans*-spliced TOP mRNAs is mediated by mTOR in *O. dioica* revealing remarkable evolutionary innovation, and flexibility, in the regulation of growth in these animals.

According to our results, exit from growth arrest in *O. dioica* appears not to be initially so responsive to nutrient levels but may instead first be triggered by a return to normal animal densities. This indicates that growth arrest in *O. dioica* exhibits some similarity to dauer arrest in *C. elegans*, which relies on the relative amounts of a dauer pheromone and a “food signal” (rather than L1 arrest, which is induced by starvation). The search for pheromomes in *O. dioica* would shed further light on this. Our observations of animals that were released from growth arrest with and without food indicate that nutrition is, however, critical for the maturation (gametogenesis) of *O. dioica* since animals that were released with food matured, spawned and died normally after 3 days whereas animals without food died after 3 days with under-developed gonads and did not spawn (data not shown). This is in agreement with our findings that the translation of *trans*-spliced TOP mRNAs is regulated by mTOR during oogenesis, as well as previous findings that egg numbers are dependent on nutrient levels (Troedsson et al., 2002): without food animals do not have sufficient resources to produce oocytes. Interestingly, animals that matured after release with food were predominantly females (81.7% females vs 18.2% males; average from 3 experiments where animals were released in the presence of food). The reason for this is unclear but we speculate that it may be due to a more flexible reallocation of resources in females according to environmental cues.

Despite sharing a common 5’ end motif in the SL, not all *trans*-spliced transcripts are affected by mTOR inhibition. We showed that many *trans*-spliced TOP mRNAs are stocked in oocytes and are translationally dormant, which largely accounts for this resistance. Interestingly, however, we also found a small subset of 21 actively translated *trans*-spliced transcripts in *O. dioica* that were resistant to Torin 1. Furthermore, in *C. elegans*, while there was an enrichment of *trans*-spliced transcripts in the set of transcripts with up-regulated translation during L1 diapause exit, a large fraction (59%; 753/1282) of transcripts with down-regulated translation are also *trans*-spliced (with SL1). This may indicate cell-type specific sequestering of a subset of SL1 transcripts upon recovery. The translation of several known TOP mRNAs is regulated in a cell-type specific manner (Avni, Biberman, & Meyuhas, 1997) and some, including poly(A) binding protein (PABP), contain downstream sequence motifs that can over-ride the TOP motif (Meyuhas & Kahan, 2015). It is possible, therefore, that mRNAs share regulatory motifs in the SL sequence but respond differently to mTOR depending on their cell-type and sequence contexts. Additional tissue-and developmental stage-specific SL-CAGE datasets would be beneficial to explore this further.

Our results show that the spliced leader in the chordate *O. dioica* is a target for nutrient-dependent translational control via mTOR. We showed that TOP mRNAs, the translation of which determines the levels of protein synthesis in the cell, are *trans*-spliced and under nutrient-dependent translational control during a developmental stage in which the majority of transcriptional output is allocated towards egg production. This supports the hypothesis that *trans*-splicing provides an evolutionary advantage in the ability to rapidly alter resource allocation during vitellogenesis according to nutritional cues from the environment. We also showed that initial recovery from growth arrest in *O. dioica* is not so responsive to nutrient levels and, accordingly, that the translational response is not dominated by *trans*-spliced transcripts. Further work on a range of additional metazoans would provide further insight into the evolution of *trans*-splicing and its role in translational control during oogenesis.

## Materials and Methods

No experiments were carried out on live vertebrates. All methods were carried out in accordance with relevant guidelines and regulations.

### Materials

Reagents were obtained from the following sources: antibodies to phospho-4E-BP1 (Thr37/46) from Cell Signalling (#2855); H3 antibodies from Abcam (#ab1791); anti-rabbit IgG secondary antibodies from KPL (#074-1506); ARTseqTM kit from Epicentre (#RPHMR12126); RNaqueous kit from Ambion (#AM1912); Torin 1 from Tocris (#4247); protease inhibitor cocktail from Sigma-Aldrich (P2714-1BTL); Halt Phosphatase Inhibitor Cocktail from Thermo Scientific (#78420); precast gels from BioRad (#4566033 and #4565013); To-Pro-3 iodide from Molecular Probes (#T3605); VECTASHIELD^®^ from Vector Laboratories (#H1000); EdU from Thermo Fisher Catalog (#A10044); Click-iT HPG Alexa Fluor 488 Protein Synthesis Assay Kit from Thermo Fisher (#C10428); TMG-cap antibodies from Santa Cruz (#sc-32724); Zymo RNA Clean & Concentrartor-25 kit (#R1018); Ribo-Zero (#MRZG12324).

### Culture of animals

Animals were cultured as previously described (Bouquet et al., 2009).

### Collection of day 6 females for ribosome profiling

At day 6, 80 female animals were transferred to 3L beakers of seawater containing regular algal food concentrations and treated with either 1 μM of Torin 1 (treatment) or DMSO (vehicle control). Animals were collected after 1.5 h and examined under a microscope to confirm they were female and to remove their houses. Collected animals were washed in PBS and frozen in liquid nitrogen.

### Collection of animals during recovery from growth arrest for ribosome profiling

Animals were collected as previously described (Danks et al., 2015) at day 7 of growth arrest and 30 minutes following release, with 700 - 1000 animals collected per sample.

### Preparation of RPF libraries

We used the ARTseqTM kit and followed the kit protocol with minor modifications. Briefly, animal samples were thawed in lysis buffer, omitting cycloheximide to avoid a 5’ end bias. Lysates were clarified by centrifugation for 10 minutes at 20,000 xg at 4°C. Amount of ARTseq nuclease was optimized, following steps in the ARTseq kit protocol, resulting in 2 units used to digest RNA. Monosomes were purified using MicroSpin S-400 columns and centrifugation for 2 minutes at 600 xg. Ribosome protected RNA fragments (RPFs) were purified using Zymo RNA Clean & Concentrator-25 kit. rRNA was removed using Ribo-Zero and RPFs PAGE purified, eluted and precipitated and resuspended in nuclease-free water and libraries prepared for sequencing following the kit protocol. Libraries were prepared in three biological replicates.

### Preparation of total RNA libraries for ribosome profiling

Total RNA was extracted from a subset of animals from each sample (15 day 6 females; 160-250 stasis/release) using the RNEasy kit and rRNA removed using Ribo-Zero. RNA was heat fragmented prior to library preparation using the ARTseq kit protocol and reagents. Libraries were prepared in three (day 6 females) or two (stasis/release) biological replicates.

### Ribosome profiling sequencing data analysis

Libraries were sequenced on two Illumina rapid flow cells at the Genomics Core Facility at the Norwegian University of Science and Technology (NTNU). We clipped adapter sequences from reads before mapping to *O. dioica* rRNA and tRNA sequences using Genoscope annotations and Bowtie2. Remaining reads were mapped to the *O. dioica* reference genome using TopHat and Genoscope gene model annotations as a guide. We used the R package *Rsubread* to calculate read counts for each protein-coding gene and the R package *babel* (Olshen et al., 2013) for differential translational efficiency analysis, with a threshold of 100 read counts across all samples for inclusion of a gene. We used previously published CAGE data mapping *SL-trans*-splice sites genome-wide (Danks et al., 2015) and classed a gene as SL *trans*-spliced if there was a *trans*-splice site within the gene body or within a 500 bp upstream region, if it was supported by >1 tag count and if it had an ‘AG’ acceptor site motif immediately upstream. Trends in translational efficiency changes were analysed by normalising all read counts to reads per million (RPM) for each library, averaging replicates and normalising RPF RPM to total RNA RPM for each gene model.

### Cap Analysis of Gene Expression (CAGE)

We extracted locations of TSSs in day 6 females from an existing CAGE data set (Danks et al., 2018). We used the sequence immediately downstream of TSSs to search for TOP and TOP-like motifs.

### Defining oocyte transcripts

We used tiling array data (Danks et al., 2013) and CAGE (Danks et al., 2018) data generated from *O. dioica* oocytes to define oocyte-stocked mRNA transcripts.

Any gene with an average probe intensity > 0 in the tiling array data or associated with a dominant CTSS ≥ 1 tpm within the gene body or 500 bp upstream region were classed as oocyte transcripts.

### Gene ontology (GO) analysis

We used *O. dioica* GO annotations (Danks et al., 2013) and the Bioconductor *GOstats* package in R to compute hypergeometric *P*-values for over-representation of GO terms.

### Collection of immature day 6 females for polysome fractionation

At day 5, female animals were separated from males, by visual inspection under a microscope, and cultured under normal conditions to day 6. At day 6, 80 immature animals were transferred to 3L beakers of seawater containing regular algal food concentrations and treated with either 1 μM of Torin 1 (treatment) or DMSO (vehicle control). Animals were collected after 1.5 h and examined under a microscope to remove their houses. Collected animals were washed in PBS and frozen in liquid nitrogen.

### Polysome fractionation

Samples were lysed with mammalian lysis buffer (200 μl 5x Mammalian polysome buffer, 100 μl 10% Triton X-100, 10μl 100mMDTT, 10 μl Dnase I (1U/μl), 678 μl Nuclease-free water) supplemented with 2 μl cyclohexamide. Lysates were placed on a sucrose gradient (15-45%) and centrifuged in a SW41 rotor at 36,000rpm for 2 hours at 4 degrees Celcius. Gradients were fractionated using the Piston Gradient Fractionator (Biocomp).

### EdU incorporation assay

Animals for growth arrest were maintained as previously. Animals were released after 7 days or food-restricted, dense culture conditions by manual transfer into clean sea water at standard culture densities, with or without standard algal strain mixture. After 12 hours, EdU (5-ethynyl-2’-deoxyuridine)-labeling assay was done according to the manual on 50 animals per sample with incubation time 30 min. Fixation in 4% PFA was followed by Click-iT reaction with Alexa 488. DNA was counterstained with 1 μM To-Pro-3 iodide and mounted in VECTASHIELD®. Samples were analyzed by confocal microscopy using a Leica TCS laser scanning confocal microscope and Leica (LAS AF v2.3) software.

### ClickIT translational assay

Click-iT HPG Alexa Fluor 488 Protein Synthesis Assay Kit was used with minor modifications. Animals were incubated in 50 μM Click-iT^®^ HPG reagent in sea water for 1 hr to label newly translated proteins. After fixation in 4% PFA, standard immunofluorescence staining (Subramaniam et al., 2014) was performed using mouse TMG-cap antibody. After secondary antibody washes, Click-iT substrate detection was performed, according to the manual, DNA was counterstained with 1 μM To-Pro-3 iodide and samples were mounted in VECTASHIELD^®^ for confocal microsopy.

### Analysis of *C. elegans trans*-splicing and ribosome profiling data

We defined *trans*-spliced genes using existing data (Allen et al., 2011). Genes with SL2 reads comprising ≤ 25% of total *trans*-spliced reads and a binomial exact P value for this percentage < 0.05 were classed as SL1. Genes with SL2 comprising ≥ 75% of reads (P value < 0.05) were classed as SL2. We classed remaining genes as “mixed” if percentages fell between these thresholds or if there were two or more *trans*-splice sites using both spliced leaders, or “unknown” if P values were ≥ 0.05.

Remaining genes (using WS207 annotations of protein coding transcripts following (Allen et al., 2011)) were classed as non-SL. WS207 gene sequence names were converted to WS230 and mapped to Wormbase protein IDs version WS235 in order to map to gene annotations used in previously generated ribosome profiling data (Stadler & Fire, 2013). We classed genes as translationally up-regulated upon exit from L1 diapause if there was a RPF fold change > 1 and an adjusted *P* value < 0.001. Genes were classed as down-regulated if there was a RPF fold change < 1 and an adjusted *P* value < 0.001. All other genes were classed as unaffected.

## Accession numbers

High-throughput sequencing data have been deposited at the NCBI Gene Expression Omnibus (http://www.ncbi.nlm.nih.gov/geo/) under accession numbers GSE78807 and GSE115265. A preprint version of this manuscript can be accessed at the following link: https://www.biorxiv.org/content/early/2018/06/22/353979.

## Author contributions

Conceptualization, G.B.D. and E.M.T.; Investigation, G.B.D., M.R., H.G., P.N., Y.T.C., and E.M.T.; Formal Analysis, G.B.D.; Visualization, G.B.D.; Validation, H.G., M.R., Y.C.T., P.N. and G.B.D.; Writing – Original Draft, G.B.D.; Writing – Review & Editing, G.B.D., H.G., M.R., P.N., E.M.T. andE.V.; Funding Acquisition, E.M.T.; Resources, E.M.T. and E.V.

## Acknowledgements

We thank Jean-Marie Bouquet, Magnus Reeve and Anne Aasjord for supplying animals as part of the animal culture facility and also for assisting in animal collections. This work was supported by grants 183690/S10 NFR-FUGE and 133335/V40 from the Norwegian Research Council (E.M.T.) High-throughput sequencing was provided by the Genomics Core Facility (GCF), Norwegian University of Science and Technology (NTNU). GCF is funded by the Faculty of Medicine at NTNU and Central Norway Regional Health Authority. The authors declare no competing interests.

## References

Allen, M., Hillier, L., Waterston, R., & Blumenthal, T. (2011, Feb). A global analysis of C. elegans transsplicing. Genome Res, 21, 255–64.

Avni, D., Biberman, Y., & Meyuhas, O. (1997, Mar). The 5’ terminal oligopyrimidine tract confers translational control on TOP mRNAs in a cell type-and sequence context-dependent manner. Nucleic Acids Res, 25, 995–1001.

Blumenthal, T., & Gleason, K. S. (2003, feb). Caenorhabditis elegans operons: form and function. Nature Reviews Genetics, 4 (2), 110–118. Retrieved from https://doi.org/10.1038%2Fnrg995 doi: 10.1038/nrg995

Bouquet, J., Spriet, E., Troedsson, C., Otterå, H., Chourrout, D., & Thompson, E. (2009, Apr). Culture optimization for the emergent zooplanktonic model organism Oikopleura dioica. J Plankton Res, 31, 359–370.

Danks, G., Campsteijn, C., Parida, M., Butcher, S., Doddapaneni, H., Fu, B., … Manak, J. (2013, Jan). OikoBase: a genomics and developmental transcriptomics resource for the urochordate Oikopleura dioica. Nucleic Acids Res, 41, D845–53.

Danks, G., Navratilova, P., Lenhard, B., & Thompson, E. (2018, Feb). Distinct core promoter codes drive transcription initiation at key developmental transitions in a marine chordate. BMC Genomics, 19, 164.

Danks, G., Raasholm, M., Campsteijn, C., Long, A., Manak, J., Lenhard, B., & Thompson, E. (2015, Mar). Trans-splicing and operons in metazoans: translational control in maternally regulated development and recovery from growth arrest. Mol Biol Evol, 32, 585–99.

Danks, G., & Thompson, E. (2015, Jul). Trans-splicing in metazoans: A link to translational control? Worm, 4, e1046030.

Douris, V., Telford, M., & Averof, M. (2010, Mar). Evidence for multiple independent origins of trans-splicing in Metazoa. Mol Biol Evol, 27, 684–93.

Ganot, P., Bouquet, J., Kallesøe, T., & Thompson, E. (2007, Feb). The Oikopleura coenocyst, a unique chordate germ cell permitting rapid, extensive modulation of oocyte production. Dev Biol, 302, 591–600.

Ganot, P., Kallesøe, T., Reinhardt, R., Chourrout, D., & Thompson, E. (2004, Sep). Spliced-leader RNA trans splicing in a chordate, Oikopleura dioica, with a compact genome. Mol Cell Biol, 24, 7795–805.

Ganot, P., Kallesøe, T., & Thompson, E. (2007, Feb). The cytoskeleton organizes germ nuclei with divergent fates and asynchronous cycles in a common cytoplasm during oogenesis in the chordate Oikopleura. Dev Biol, 302, 577–90.

Ganot, P., Moosmann-Schulmeister, A., & Thompson, E. (2008, Dec). Oocyte selection is concurrent with meiosis resumption in the coenocystic oogenesis of Oikopleura. Dev Biol, 324, 266–76.

Hastings, K. (2005, Apr). SL trans-splicing: easy come or easy go? Trends Genet, 21, 240–7.

Hsieh, A., Liu, Y., Edlind, M., Ingolia, N., Janes, M., Sher, A., … Ruggero, D. (2012, Feb). The translational landscape of mTOR signalling steers cancer initiation and metastasis. Nature, 485, 55–61.

Ingolia, N., Brar, G., Rouskin, S., McGeachy, A., & Weissman, J. (2012, Jul). The ribosome profiling strategy for monitoring translation in vivo by deep sequencing of ribosome-protected mRNA fragments. Nat Protoc, 7, 1534–50.

Krchňáková, Z., Krajčovič, J., & Vesteg, M. (2017, jul). On the Possibility of an Early Evolutionary Origin for the Spliced Leader Trans-Splicing. Journal of Molecular Evolution, 85(1-2), 37–45. Retrieved from https://doi.org/10.1007%2Fs00239-017-9803-y doi: 10.1007/s00239-017-9803-y

Levy, S., Avni, D., Hariharan, N., Perry, R., & Meyuhas, O. (1991, Apr). Oligopyrimidine tract at the 5’ end of mammalian ribosomal protein mRNAs is required for their translational control. Proc Natl Acad Sci USA, 88, 3319–23.

Meyuhas, O. (2001, December). Synthesis of the translational apparatus is regulated at the translational level. European Journal of Biochemistry, 267(21), 6321–6330.

Meyuhas, O., & Kahan, T. (2015, Jul). The race to decipher the top secrets of TOP mRNAs. Biochim Biophys Acta, 1849, 801–11.

Olshen, A., Hsieh, A., Stumpf, C., Olshen, R., Ruggero, D., & Taylor, B. (2013, Dec). Assessing gene-level translational control from ribosome profiling. Bioinformatics, 29, 2995-3002.

Stadler, M., & Fire, A. (2013). Conserved translatome remodeling in nematode species executing a shared developmental transition. PLoS Genet, 9, e1003739.

Subramaniam, G., Campsteijn, C., & Thompson, E. (2014). Lifespan extension in a semelparous chordate occurs via developmental growth arrest just prior to meiotic entry. PLoS One, 9, e93787.

Thoreen, C., Chantranupong, L., Keys, H., Wang, T., Gray, N., & Sabatini, D. (2012, May). A unifying model for mTORC1-mediated regulation of mRNA translation. Nature, 485, 109–13.

Thoreen, C., Kang, S., Chang, J., Liu, Q., Zhang, J., Gao, Y., … Gray, N. (2009, Mar). An ATP-competitive mammalian target of rapamycin inhibitor reveals rapamycin-resistant functions of mTORC1. J Biol Chem, 284, 8023–32.

Troedsson, C., Bouquet, J. M., Aksnes, D., & Thompson, E. M. (2002, November). Resource allocation between somatic growth and reproductive output in the pelagic chordate Oikopleura dioica allows opportunistic response to nutritional variation. Marine Ecology Progress Series, 243, 83–91.

Yang, Y., Zhang, X., Ma, X., Zhao, T., Sun, Q., Huan, Q., … Qian, W. (2017, Sep). Trans-splicing enhances translational efficiency in C. elegans. Genome Res, 27, 1525-1535.

Yokomori, R., Shimai, K., Nishitsuji, K., Suzuki, Y., Kusakabe, T., & Nakai, K. (2016, Jan). Genome-wide identification and characterization of transcription start sites and promoters in the tunicate Ciona intestinalis. Genome Res, 26, 140–50.

Zaslaver, A., Baugh, L., & Sternberg, P. (2011, Jun). Metazoan operons accelerate recovery from growth-arrested states. Cell, 145, 981–92.

