## Supplemental figures for "Trans-splicing of mRNAs links gene transcription to translational control regulated by mTOR"

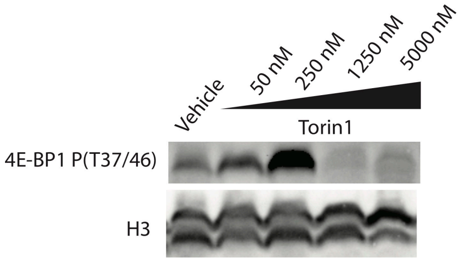


**Figure S1. Dosage response of *O. dioica* to increasing concentrations of the mTOR inhibitor Torin 1.**
Female animals were exposed to DMSO (vehicle control) and different concentrations (50 nM, 250 nM, 1250 nM and 5000 nM) of the mTOR inhibitor, Torin 1, and 4E-BP1 phosphorylation levels were assayed. Histone H3 was used as a reference loading control.

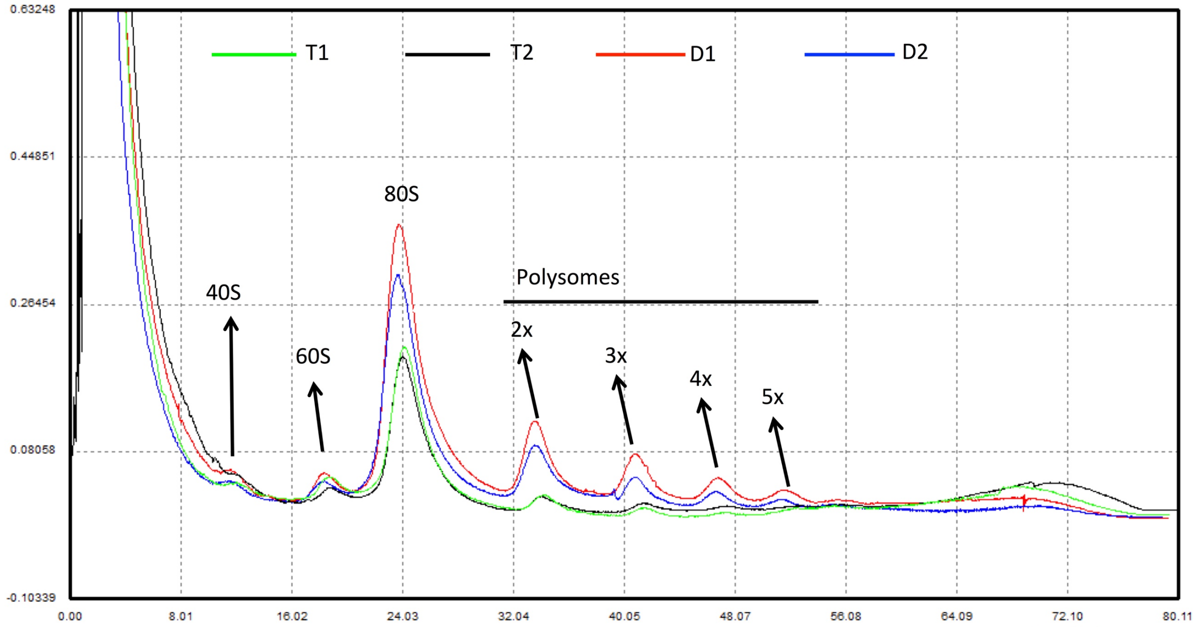


**Figure S2. Translational response to treatment with the mTOR inhibitor Torin1.**
Polysome profiles from two replicates of treated (T1 and T2) and control (D1 and D2) day 6 animals confirmed a down-regulation of translation in treated animals as indicated by reduced polysome peaks.

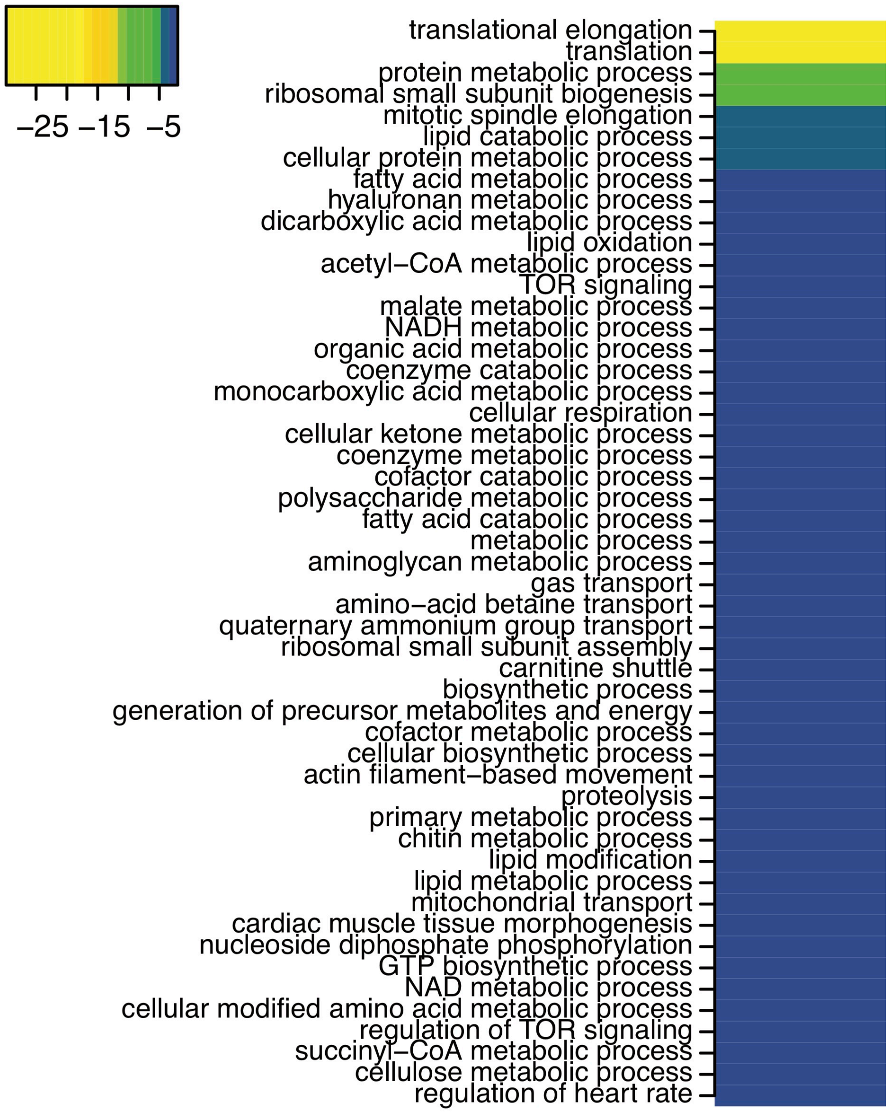


**Figure S3. Conserved functions in the targets of mTOR-dependent translational control in *O. dioica*.**
GO terms and p-values from a gene ontology analysis of genes with transcripts that were significantly down-regulated translation upon mTOR inhibition with Torin 1.

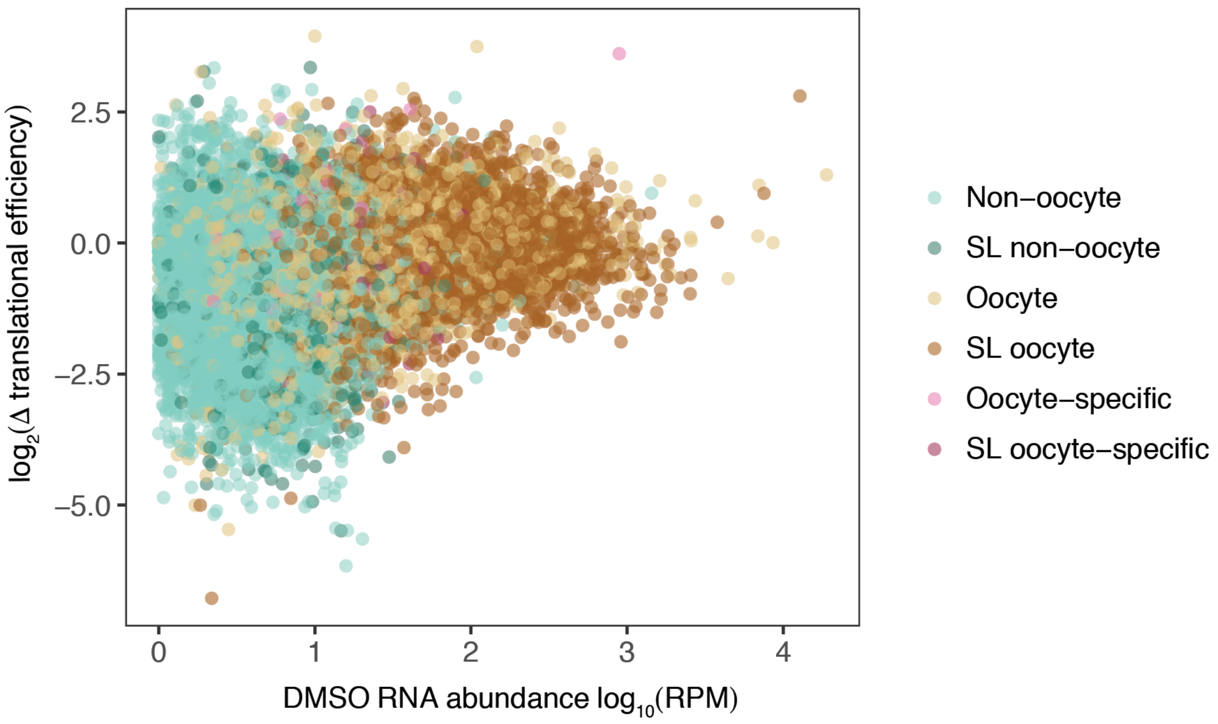


**Figure S4. Oocyte transcripts are *trans*-spliced and translationally dormant.** Changes in translational efficiency in response to Torin 1 (y axis) against mRNA abundances (RPM = reads per million) in control animals (x axis) with transcripts categorised as indicated in the legend.

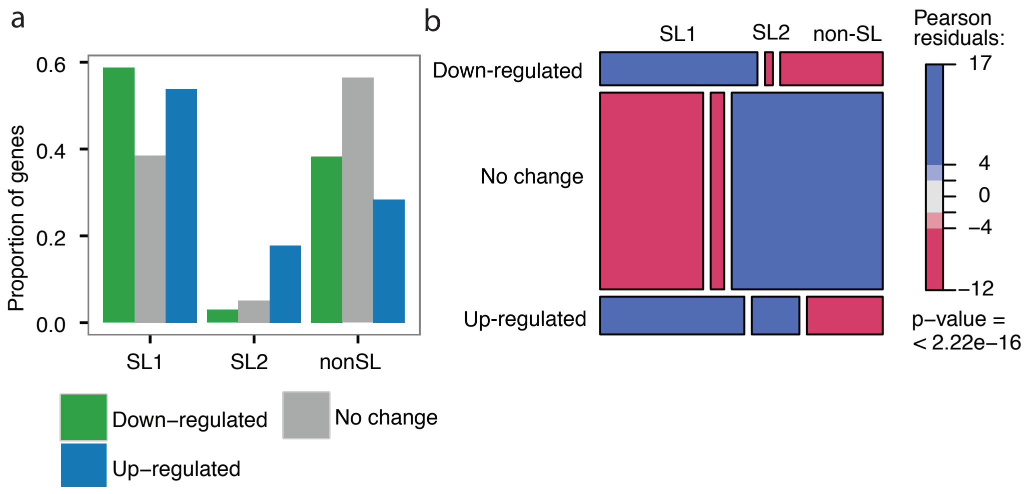


**Figure S5. Translational control during nutrient-dependent recovery from growth arrest is associated with the presence of a 5' spliced leader in *C. elegans*.** (A) Proportion of genes *trans*-spliced to SL1 or SL2 or without a spliced leader that have translation up- or down-regulated (or no translational response) upon release from L1 diapause in response to food availability. (B) Mosaic plot shows Pearson residuals from a Chi-square test using genes categorised as in (A).

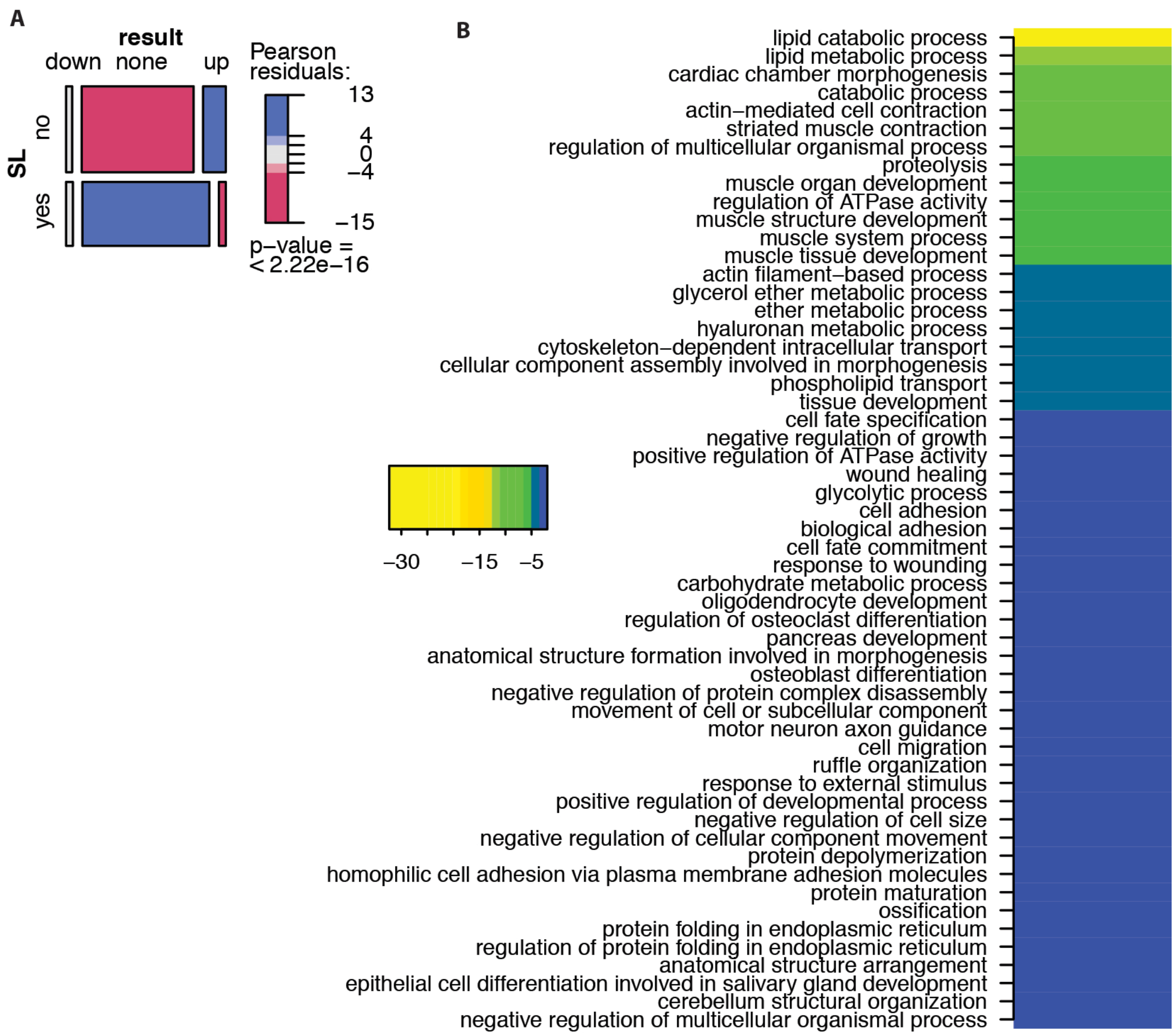


**Figure S6. Transcriptional response during recovery from growth arrest in O. dioica.**

Genes with significantly up-regulated transcription during recovery from growth arrest were enriched for non-*trans*-spliced transcripts (A) and GO terms related to lipid metabolism, muscle contraction and proteolysis (B).

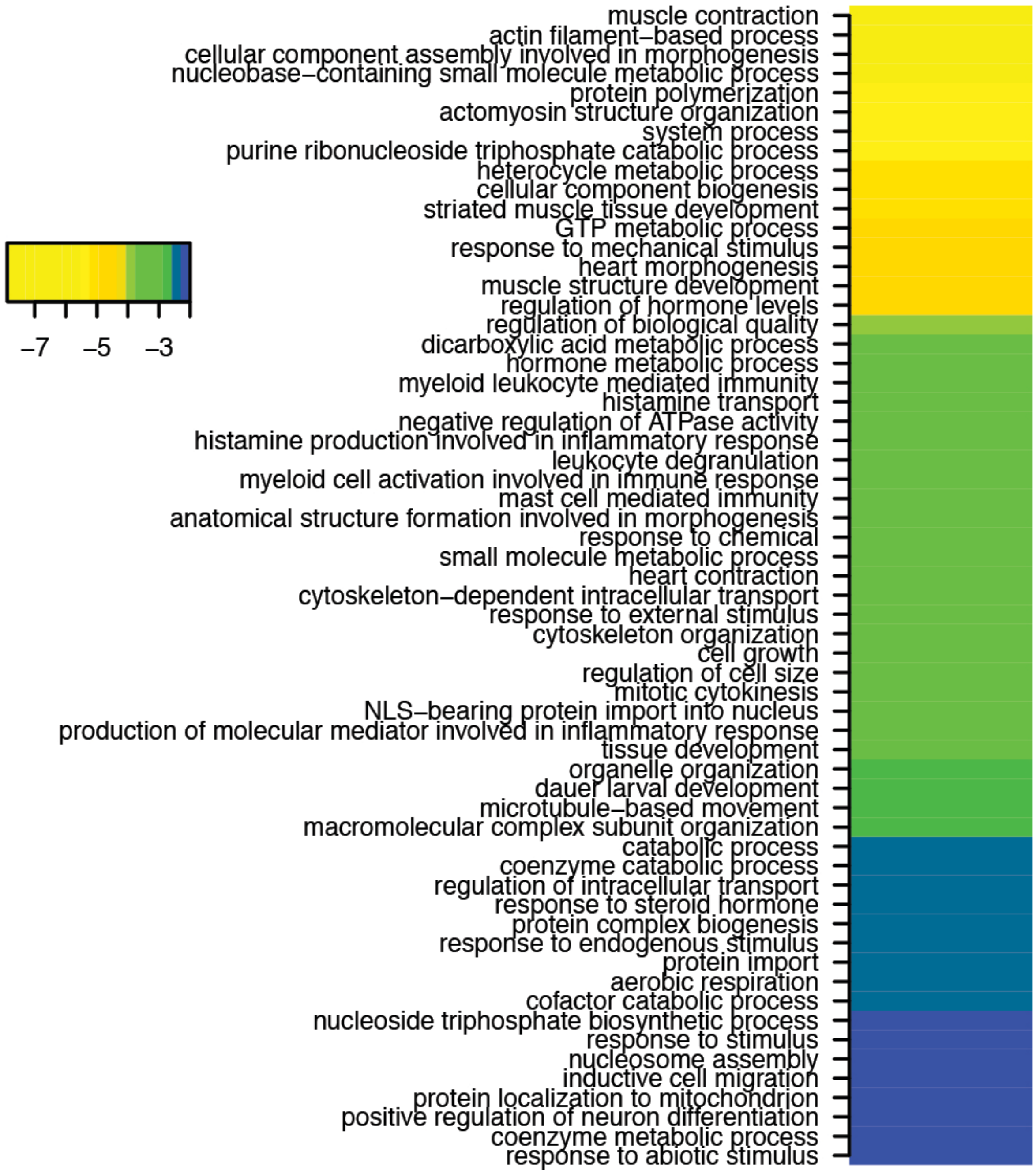


**Figure S7. Translational response during recovery from growth arrest in O. dioica.**

GO terms enriched in genes with significantly up-regulated translation during recovery from growth arrest.
